# Non-invasive detection of local microstructural damage in tendon via Diffusion Tensor MRI

**DOI:** 10.1101/2025.10.10.681653

**Authors:** Roberto A. Pineda Guzman, Amir Ostadi Moghaddam, Matthew P. Confer, Shreyan Majumdar, Rohit Bhargava, Amy J. Wagoner Johnson, Bruce M. Damon, Mariana E. Kersh

## Abstract

Tendon is critical for musculoskeletal function as it transfers forces generated by muscle to bone and stores energy during movement. Impaired mechanical function in tendon limits mobility and results from fatigue-induced damage progression that outpaces the restorative processes maintaning tissue health, a phenomenom we term *mechanopathology*. Early and non-invasive detection of tendon mechanopathologies is vital to prevent further damage, but is lacking in the clinical space. Here, we evaluate the ability of diffusion tensor magnetic resonance imaging (DT-MRI) to detect mechanical fatigue damage in tendon, and validate our findings using histologic assessments of collagen fiber microstructure and molecular structure. We found that fatigue-induced changes in DT-MRI metrics of tendon are spatially heterogeneous, and correspond to regions with damaged collagen fiber microstructure. While secondary structures of collagen molecules were damaged by fatigue loading, they do not spatially correspond to fatigue-induced changes in DT-MRI metrics. Fatigue-induced changes in DT-MRI metrics can be partially explained by quantitative metrics of post-fatigue collagen fiber microstructure, estimating the limit of detection of DT-MRI metrics to fatigue-induced damage in tendon. Our findings indicate that DT-MRI metrics are sensitive to fatigue-induced local damage in tendon, supporting the potential of DT-MRI as a non-invasive and translatable tool to clinically detect mechanopathologies in tendon.

## 1. Introduction

Musculoskeletal conditions are among the leading causes of years living with disability, contributing to diminished quality of life, and yet progress towards their early detection remains elusive [1]. Tendon is critical for musculoskeletal function as it serves to transfer forces generated by muscles to bone and stores energy during movement. The hierarchal organization of tendon provides resilience and enables a lifetime of loading cycles in healthy individuals. However, impaired mechanical function in tendon limits mobility, and results from microstructural damage progression that outpaces restorative processes that maintain tissue health [2, 3], a phenomenon we term *mechanopathology*. Mechanopathologies are more frequently observed in physically active individuals [4, 5, 6] as well as those whom engage in non-normative loading such as wheelchair users [7]. In both cases, repeated loading originates fatigue-induced damage progression and ultimately leads to bulk tissue failure [8]. Non-invasive assessment of mechanopathologies is vital to identifying tendon damage prior to failure [2, 9, 10], especially at an early stage, but are unfortunately lacking in the clinical space.

The diagnosis of tendon mechanopathologies is complicated by the lack of early symptoms [11], the highly variable individual response to fatigue loading [12, 13, 14], and limitations in the methods used for non-invasively detecting the effects of fatigue loading. Ultrasound, for example, is the primary imaging modality used to detect tendon damage [15, 16, 17, 18, 19, 20], but has limited field of view and is constrained to imaging superficial tendons [21]. Detecting fatigue-induced damage in tendon requires whole-tissue assessments, as *in vitro* studies have reported significant spatial heterogeneity in the mechanical [22] and microstructural [23] response of tendon to fatigue loading. The local detection of tendon damage is possible using histological methods [24, 8, 25], but suffer from similar challenges of sparse spatial sampling capabilities and are typically infeasible in the clinical setting. Thus, full-field and non-invasive imaging techniques that characterize microstructural damage are needed to assess tendon damage prior to failure *in vivo*.

Magnetic Resonance Imaging (MRI) is routinely used to diagnose musculoskeletal injuries based on qualitative assessments of whole tissue structure [26], but has unfulfilled translational potential to *quantitatively* assess tissue microstructure. Diffusion Tensor Magnetic Resonance Imaging (DT-MRI) is sensitive to tissue microstructure, as it measures hindrances in water diffusion caused by the presence of physical barriers within the tissue [27]. The magnitude of diffusion in an anisotropic fiber-based tissue is typically characterized using three metrics: 1) Axial Diffusivity (AD) measures diffusion along the preferred orientation of the physical barriers, 2) Radial Diffusivity (RD) which measures diffusion perpendicular to the physical barriers, and 3) Apparent Diffusion Coefficient (ADC) measures the average diffusion along three spatial dimensions. DT-MRI metrics are widely used to study brain microstructure [28, 29, 30] and, to a lesser extent, skeletal muscle [31, 32, 33]. Despite technical challenges associated with low signal-to-noise ratio (SNR) of DT-MRI data in tendon and ligament [34, 35, 36, 37], advancements in hardware and image acquisition schemes have established the feasibility of DT-MRI in these tissues [34, 38, 39, 40, 41, 42, 43]. For example, DT-MRI metrics have been used in longitudinal evaluations to monitor ligament healing following surgical interventions [44, 45]. However, the mechanopathological interpretation of DT-MRI metrics and their relationship to damage are still not understood. To our knowledge, no previous study has assessed the effect of fatigue-induced damage on DT-MRI metrics of connective soft tissues.

The early detection of mechanopathologies using DT-MRI metrics requires evaluating their sensitivity to mechanically-induced microstructural and molecular damage [46]. Collagen - the fundamental building block of soft musculoskeletal tissues, including tendon - is a structural protein that forms into fibrillar structures and plays a vital role in musculoskeletal tissue function [47]. Previously, we found that fatigue-induced damage in tissue-mimicking fibers results in an increase in ADC, AD, and RD measured in the interfibrillar water [48]. What remains is to validate these findings in ligament and tendon, tissues with heterogeneous collagen microstructure and other extracellular matrix components, using ground-truth histological measurements of collagen fiber microstructure [49, 50] that are sensitive to fatigue-induced damage [51, 23]. Moreover, given that the hierarchical organization of tendon contributes to its evolving mechanical properties at each length scale, the response of molecular-level assemblies to fatigue-induced damage, and their effect on the DT-MRI metrics of tendon, warrants attention but has been underexplored[23, 52].

In this study, we demonstrate that DT-MRI metrics are altered by *in vitro* mechanical fatigue damage in tendon. We show that the fatigue-induced changes in DT-MRI metrics are spatially heterogeneous and are observed in areas with damaged collagen fiber microstructure, quantified via Second Harmonic Generation (SHG) microscopy. Using Fourier-transform Infrared (FTIR) spectroscopy, we show that secondary molecular structures are damaged by fatigue loading but do not spatially correspond to fatigue-induced changes in DT-MRI metrics. Finally, we establish the proportion of changes in DT-MRI metrics that could be explained by quantitative metrics of post-fatigue collagen fiber microstructure, thereby providing an estimate of the limit of detection of 9.4 tesla-based DT-MRI metrics to fatigue-induced damage. Collectively, these analyses confirm the sensitivity of DT-MRI metrics to fatigue-induced local damage in tendon, and the degree to which changes in DT-MRI metrics are related to changes in collagen fiber microstructure and molecular integrity, thus supporting the use of DT-MRI metrics as a non-invasive biomarker of mechanopathologies in tendon.

## 2. Materials and Methods

### 2.1. Study design and summary of approach

The goal of this study was to evaluate the ability of DT-MRI metrics to detect fatigue-induced damage in tendons *in vivo*, and relating the observed changes in DT-MRI metrics to damage observed in the underlying collagen fiber microstructure and molecular structure of fatigued tendons.

The DT-MRI fatigue experiment consisted in subjecting a group of tendons (*n* = 5) to the following protocol: 1) an initial DT-MRI scan, 2) a fatigue loading protocol after the first scan, and 3) a second DT-MRI scan of the tendon after being fatigued (Figure 1A). The differences in DT-MRI metrics between the pre and post fatigue scans were quantified to describe fatigue-induced changes. To account for changes in DT-MRI metrics that could be caused by tissue decomposition between scans, a group of control tendons (*n* = 5) underwent the following protocol: 1) an initial DT-MRI scan, 2) a control protocol that replicated all the experimental conditions of the fatigued samples except for the fatigue loading, and 3) a second DT-MRI scan of the tendon.

**Figure 1.**
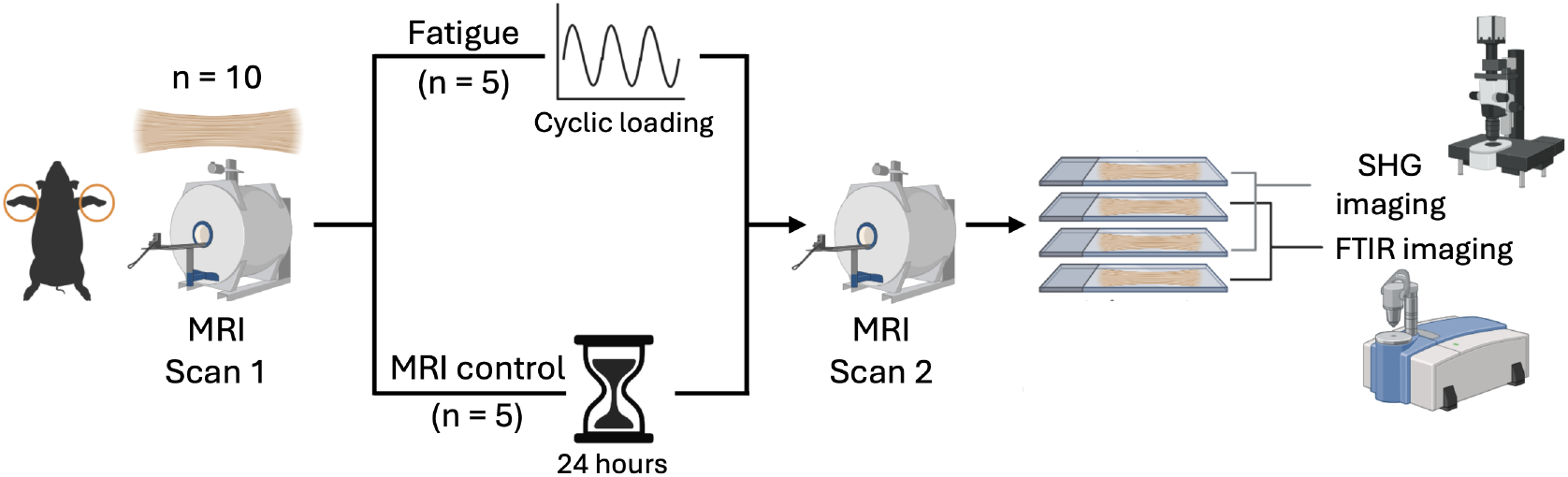
Study design: schematic of experimental setup for detection of fatigue-induced microstructural damage in tendon using DT-MRI. Two groups of tendons (control and fatigue) were studied to assess the effect of fatigue-induced damage on the DT-MRI metrics of tendon. The DT-MRI metrics of the tendons were measured before and after undergoing cyclic loading or control conditions. To assess the role of microstructural and molecular determinants in the observed changes in DT-MRI metrics, SHG and FTIR data were collected from the tendon samples following the second DT-MRI scan.

To analyze the underlying collagen fiber microstructure and molecular composition of both fatigue and control tendons, co-registered SHG and FTIR images of the tendons were obtained after the second DT-MRI scan from interleaved microscopic slices (Figure 1B). To account for spatial heterogeneity within the tendon volume, all tendons were divided in nine distinct regions where DT-MRI, SHG, and FTIR metrics were assessed (Figure 1C). The nine regions were obtained from three subdivisions across the length of the tendon and three radial subdivisions across the tendon’s cross-section, similar to a previous study in deep digital flexor tendons [53]. Regional changes in the DT-MRI metrics of the tendons caused by the fatigue and control experiments were computed. Differences between the SHG and FTIR metrics of fatigue and control tendons were quantified to characterize fatigue-induced microstructural and molecular changes. The correlations between the regional changes in DT-MRI metrics and the SHG metrics in the fatigued group were evaluated to establish the relationship between spatial differences in fatigue-induced collagen fiber microstructure and the measured changes in the tendon DT-MRI metrics. No previously published data of the DT-MRI metrics of damaged and non-damaged tendon is available, and a preliminary power analysis could not be conducted prior to the experiments.

### 2.2. Sample preparation

Ten porcine feet were acquired from a local grocer and stored at -20*°*C. The feet were thawed at 4*°*C between 36 and 48 hours prior to dissection. The pair of deep digital flexor tendons was harvested from each foot by cutting both tendons on their bone insertion and the muscle belly with a number 20 scalpel blade. The muscle tissue was carefully removed from the musculotendinous junction and the two deep digital flexor tendons were separated at their anatomical bifurcation point using a number 10 scalpel blade. One of the two tendons was randomly selected and cut into a 100 mm long section that included 30 mm from the tendon’s bifurcation and 70 mm from the tendon midsubstance. A tissue marking dye (MarkIt, Thermo Fisher Scientific, MA, USA) was used to mark the proximal edge of a 20 mm-long central area of the tendon section. This mark was used to help ensure that the tendons were imaged in the same locations in the two DT-MRI scans and the SHG and FTIR images.

### 2.3. Initial DT-MRI scan

The sample preparation procedure followed prior to the DT-MRI scan is described in the supplementary materials (Figure S2A). The tendons from the fatigue and control groups (*n* = 5 for each group) were scanned in a 9.4 tesla MRI scanner (Bruker, MA, USA) using a circularly polarized volume coil with an outer/inner diameter of 112/86 mm for transmission, and a ^1^H receive-only 2 × 2 rat brain array coil for reception. DT-MRI was performed using a diffusion-weighted spin echo sequence (Echo time = 18 ms, Repetition time = 800 ms, 12 diffusion encoding directions, 15 averages, b-value = 400 s/mm^2^, diffusion gradient duration = 4 ms, diffusion gradient separation = 10 ms, and scan time = 2.6 hours). Ten slices were acquired, with a slice thickness of 2 mm, an in-plane resolution of 0.5 × 0.5 mm^2^, and a field of view of 20 × 20 × 20 mm^3^. A registration marker filled with MRI visible fluid was placed over the proximal edge of the tissue dye mark, setting the edge of the scan’s field of view (Figure S2B). The samples were then stored inside a centrifuge tube overnight at 4*°*C in a refrigerator.

### 2.4. Fatigue loading and post-fatigue DT-MRI scan

The tendons in the fatigue group (*n* = 5) underwent a fatigue loading protocol the day after the initial DT-MRI scan using a uniaxial tensile testing system (Instron 5967, Instron, MA, USA). The 30 mm proximal and distal ends of the tendon samples were attached to sandpaper tabs using cyanoacrylate glue. The tendon and sandpaper tabs were then clamped with serrated grips that were connected to a pneumatic actuation system at a pressure of 600 kPa. The grip-to-grip testing length was 40 mm, with the 20 mm imaging region centered between the grips (Figure 1B). Preliminary tensile tests were conducted in four deep digital flexor tendon samples to obtain an estimate of the ultimate tensile load of the tendons. The preliminary testing samples were ramped to failure at a strain rate of 0.5%/s, and an ultimate tensile load of 683 *±* 67 N was measured.

The tendons from the fatigue group were subjected to a displacement-controlled loading protocol specified between 100 and 273.2 N (40% of the tendon’s ultimate load), at a strain rate of 0.6%/s for 10,000 loading cycles. Fatigue loading data were collected at 100 Hz. The fatigue protocol strain rate was set to obtain a loading frequency within 1-2 Hz. The number of loading cycles was chosen to create accumulated damage prior to failure, which is expected at 128882 loading cycles based on a previous study on human Achilles tendon [12]. The tendons were kept moist during fatigue testing with two phosphate buffered saline (PBS) drips positioned in the proximal edge of the tendon with a flow rate of 20 mL/hour and by spraying PBS over the whole tendon surface every 10 minutes.

The same section of the tendons scanned in the pre-fatigue scan was scanned post-fatigue, using the same diffusion-weighted spin echo sequence. The steps followed to ensured that the same area of the tendon was assessed in the pre and post-fatigue DT-MRI scans are described in the supplementary materials (Figure S2C-D).

### 2.5. Control experiment and Day 2 DT-MRI scan

The tendons in the control group (*n* = 5) were taken out of the refrigerator the day after the initial DT-MRI scan. Instead of undergoing fatigue loading, the tendons were left inside the centrifuge tube at room temperature for 2 hours, the approximate duration of the fatigue loading protocol. The same section of the tendons scanned in the initial DT-MRI scan was scanned in the second scan, using the same diffusion-weighted spin echo sequence used in the first scan. The field of view of the second scan was located using the tissue dye mark placed on the tendon, ensuring that the same area of the tendon was imaged in both DT-MRI scans. After the second DT-MRI scan, the tendon samples were frozen at -20*°*C.

### 2.6. Second harmonic generation microscopy

Second-harmonic generation (SHG) imaging was carried out on the cryosectioned tendons using a Zeiss LSM 710 multiphoton microscope (Zeiss, Oberkochen, Germany) equipped with a MaiTai DeepSee Ti:Sapphire femtosecond laser (Spectra-Physics, Newport Corp., USA) tuned to 780 nm (70 fs pulses). Samples were thawed in 1× PBS for 10 minutes, and a tissue dye mark was reapplied to the proximal tendon edge to preserve orientation. From this mark, 22 mm segments were excised: 20 mm of each segment encompassed the region of interest, with 1 mm buffers at both ends. Segments were embedded in OCT such that the tendon’s long axis lay horizontally, with the marker positioned at the lower right of the mold. Sequential 50 µm cryosections were collected at 500 µm intervals onto No. 1.5 glass slides maintained at –20 *°*C, while intervening 5 µm sections were reserved for FTIR analyses. Immediately before imaging, each 50 µm section was thawed, rehydrated with 1–2 drops of PBS, and gently covered with a coverslip—lowered slowly using fine-tip forceps to avoid air bubbles. SHG images were acquired with a 10×/0.30 NA air objective at 1× digital zoom, producing 512 × 512 pixels per tile (1.66 µm/pixel calibration). A pixel dwell time of 1.272 µs and line period of 30 µs were employed without line averaging. Tiles were automatically connected in Zen software (Zeiss). Emission was collected in both forward and backward directions, with laser power and detector gain held constant across all sections.

### 2.7. FTIR microscopy

Five *µ*m thick tendon samples were cryosectioned on IR reflective low-emissivity slides (Kevley Technologies, OH, USA). These sections were stored at -80*°*C until use. Samples were then thawed for 20 minutes under low vacuum prior to fixation with methanol-free 4% paraformaldehyde and quenching with glycine. The samples were then allowed to dry under N_2_ flow prior to imaging. FTIR microscopy was performed using an Agilent 870 imaging system (Agilent, CA, USA) equipped with a 128 × 128 element focal plane array Mercury-Cadmium-Telluride (MCT) detector. The images were acquired at 5.5 *µ*m pixel size with 4 sample coadditions and 128 background coadditions. Spectroscopic images were collected at 4 cm^*−*1^ spectral resolution and 2 cm^*−*1^ spacing from 800-3950 cm^*−*1^.

### 2.8. Data analysis

#### 2.8.1. Fatigue loading data

The maximum cyclic displacement, *u*_*max,n*_, was computed as the displacement in which the maximum force was measured in cycle (*n*). The cyclic stiffness was computed as:

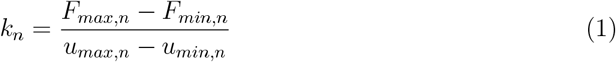

where *F*_*max,n*_ and *F*_*min,n*_ are the maximum and minimum forces measured in cycle (*n*), and *u*_*min,n*_ is the displacement in which *F*_*min,n*_ was measured. *u*_*max,n*_ and *k*_*n*_ were measured in cycles 1 and 5000. Sample 3 had a failed initial loading protocol where it underwent 3089 loading cycles with considerable sample sliding. Because sliding introduces inaccuracies to the displacement data from the initial fatigue loading test, the data reported in sample 3 corresponds to a second loading test initiated after re-gripping the sample. In this new fatigue loading test, cycles 1 and 5000 correspond to the cycles 3090 and 8089 applied to sample 3.

#### 2.8.2. DT-MRI data

The DT-MRI data of one sample from the control group was excluded due to motion artifacts caused by a malfunction in the scanner’s bed. The diffusion-weighted spin echo images from the remaining samples showed no observable artifacts or distortion in the non-diffusion weighted and diffusion weighted images, and did not require image registration as a pre-processing step. All DT-MRI data analyses were conducted in MATLAB R2023b (MathWorks, MA, USA). Tendon masks were created from the non-diffusion weighted (b = 0) images by segmenting all voxels with a 75% decrease in the signal relative to the signal measured in the PBS surrounding the tendon. Signal-to-noise ratio (SNR) estimates of the non-diffusion weighted images of the tendons were estimated by dividing the mean signal intensity in the tendon by a noise estimate equal to 1.25 times the mean signal in an empty section at the top-left corner of the image [54]. SNR estimates were then corrected to account for the bias present in magnitude images obtained from phase arrray systems using sum-of-squares reconstruction [55]. The mean SNR computed in all the tendon scans was 21.0 *±* 4.35 (range, 14.8 - 31.1). This SNR level is insufficient to obtain accurate DT-MRI metrics of anisotropic structures at the voxel level [35, 36]. To reduce the bias caused by the low SNR measured in the tendon, we grouped the tendon’s voxels in nine different regions to implement a signal averaging strategy that improves the accuracy of the DT-MRI metrics [35, 48] and accounts for the microstructural heterogeneity along the tendon’s anatomy. The nine regions were defined by three radial and three longitudinal regions along the tendon. The centroid of the tendon’s cross section in each slice was computed and the voxels with a radius lower than 1 mm were included in the central region, those with a radius higher than 1 mm and lower than 2 mm were included in the intermediate region, and those with a radius higher than 2 mm were included in the outer region. Longitudinally, the tendons were divided into three different regions: a proximal region consisting of the three most proximal slices of the tendon MRI image, a middle region consisting of the four central slices of the tendon MRI image, and a distal region consisting of the three most distal slices of the tendon MRI image (Figure 1C).

The average signals measured in the non-diffusion and diffusion weighted images of each tendon region were used to compute the region’s diffusion tensor using a weighted linear-least squares algorithm. This signal averaging method reduces the bias of low SNR [35], therefore increasing the accuracy of the measured DT-MRI metrics. The diffusion tensor was diagonalized using singular value decomposition to compute the tensor eigenvalues (*λ*_1_, *λ*_2_, and *λ*_3_). The Apparent Diffusion Coefficient (ADC) was calculated as:

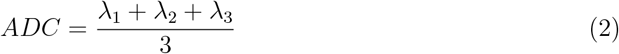

while Axial Diffusivity (AD) was computed as the first eigenvalue (*λ*_1_), and Radial Diffusivity was computed as the mean of *λ*_2_ and *λ*_3_. We ran a sensitivity study to ensure that ADC, AD, and RD were accurately measured at the lowest SNR level measured in each radial region, while also finding that other DT-MRI metrics such as fractional anisotropy and the first eigenvector of the diffusion tensor can not be accurately measured using our signal averaging strategy. Detailed information of the sensitivity study is shown in the supplementary materials [56].

#### 2.8.3. SHG data

SHG image stacks were analyzed in MATLAB (R2024a) using a custom Fourier-transform SHG (FT-SHG) pipeline. Each stack was down-sampled to achieve isotropic voxels (0.5 µm pitch) and then subdivided into cubic sub-volumes (6 × 6 × 6 µm) chosen based on collagen fiber diameter and crimp wavelength to guarantee adequate orientational resolution. For every sub-volume the Fourier power spectrum was computed; the principal fiber direction was taken as the orientation of the maximum in this spectrum, and its magnitude provided a normalized weight that accounts for local anisotropy. Select regional data was discarded due to lack of SHG signal from the region of interest.

From these data we derived three texture metrics—% Dark, % Isotropic and circular variance (CV)—which were averaged over each radial region. % Dark represents the fraction of voxels whose mean SHG intensity falls below a fixed threshold set to 10% of the global maximum image intensity, identifying collagen-poor or fibers oriented parallel to the optical axis. % Isotropic denotes voxels lacking a clear spectral peak, indicative of uniformly dispersed or non-fibrillar regions. CV quantifies the in-plane angular dispersion on a continuum from 0 (perfect alignment) to 1 (complete disorder), following the conventions of our earlier work [57, 58, 59].

To link SHG signal intensity directly to diffusion-MRI descriptors we calculated L2, an eigenvalue analogue. Within each 2 mm × 0.5 mm tile, fiber orientations were re-sampled on a 32 × 32 pixel grid; dark and isotropic cells were discarded, yielding *n* anisotropic unit vectors *u*_*i*_ = (*d*_*x,i*_, *d*_*y,i*_). Vectors were sign-adjusted so that all lay within *±*90° of the tile-wide mean axis, enforcing axial symmetry. We then formed the in-plane orientation tensor

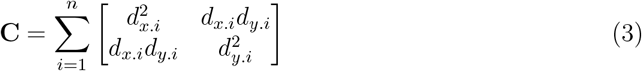

and diagonalized it to obtain eigenvalues *λ*_1_ ≥ *λ*_2_. Treating this result as a diffusion-like 3 × 3 tensor (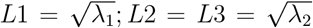), we defined the scalar L2 to quantify residual orientation variance orthogonal to the dominant fiber direction: L2 approaches 0 for highly aligned fibers and approaches L1 as orientation becomes azimuthally isotropic. The average L2 was computed in each radial region.

#### 2.8.4. FTIR data

Image processing was performed using a combination of custom Python and Matlab 2024a (MathWorks, MA, USA) codes and ENVI (NV5 Geospatial Solutions, FL, USA). Image tiles were connected before noise reduction was performed using a minimum noise fraction transform. Spectra were baseline corrected to account for scattering and normalized to the amide I peak. Each tendon was then masked to remove background pixels and regions where the tissue had lifted off the microscope slide. Tendons were divided into nine regions for further analysis (Figure 1C). Radial subdivisions were divided by determining the outline of the sample, fitting a second order polynomial function through the long axis of the tendon, and then offsetting this polynomial in the radial direction. Longitudinal subdivisions were determined by dividing the sample along the normal lines to the second order polynomial used for radial region division [60]. The tendon sections used for FTIR microscopy did not contain sufficient data in the outer radial (R3) region to allow for proper analysis, therefore only the central and intermediate radial regions were analyzed.

### 2.9. Statistical analysis

Statistical analyses were conducted using R (version 4.3.3). Outliers were identified using the Grubbs test and were removed from the regional datasets of each DT-MRI and SHG metric. FTIR data quality was inspected instead of outlier removal, and no data points were removed. After removing outliers, statistical analyses were conducted separately on the DT-MRI, SHG, and FTIR data separately. Visual inspections of quantile-quantile plots generated by the *ggqqplot* function were made to determine the normality of the data distributions and determine the need of parametric or non-parametric statistical tests. For the DT-MRI metrics, 2-way repeated measures Aligned Ranked Transform Analysis of Variance (ART-ANOVA) [61] tests were used to evaluate the effect of fatigue loading on the DT-MRI metrics of the tendons, considering the tendon samples as a random effect and accounting for the interaction between anatomical region and fatigue response. For the SHG metrics, were there was missing data in some anatomical regions, the global differences between the control and fatigue group were evaluated considering the tendon samples as a random effect and accounting for the effect of fatigue without the interaction of anatomical regions. Analysis of Variance (ANOVA) was used to evaluate the differences in CV, % Isotropic, and L2 while ART-ANOVA was used to evaluate % Dark. Tests evaluating the differences in FTIR metrics between the control and fatigue group were conducted by considering the tendon samples as a random effect and accounting for the interaction between fatigue loading and anatomical region. A 2-way ART-ANOVA was conducted for all the FTIR metrics, except for the absorbance at 1558 cm^*−*1^, in which a 2-way ANOVA was used. Regional differences in SHG and FTIR metrics were evaluated separately. Normality of regional datasets was evaluated using a Shapiro-Wilks test and differences between the fatigue and control group were evaluated with unpaired t-tests or ks-tests. Holm-Bonferroni corrections were applied to control for multiple comparisons. Correlations between the changes in DT-MRI metrics and SHG metrics were made in the fatigued and control group using repeated measure correlations [62], considering the regional measurements made in each tendon sample as repeated measurements.

## 3. Results

### 3.1. Moderate fatigue results in measurable mechanical changes in tendon

To quantify the effect of mechanical fatigue on tendon microstructure using DT-MRI metrics, we performed a series of fatigue experiments in flexor tendons (n=10 samples) with DT-MRI data collected before and after fatigue loading (Figure 1). We mechanically damaged a group of tendon samples (fatigue group, *n* = 5) using a cyclic tensile loading protocol. While a key strength of MRI is the capacity to perform longitudinal within sample measurements, we included a control group to ensure that the detected damage in the fatigued tendons was not caused by experimental conditions unrelated to the fatigue loading protocol. Therefore, the remaining five samples (control group, *n* = 5) replicated all experimental conditions, including changes in temperature and timeline, of the tendons in the fatigued group but did not undergo a fatigue loading protocol.

All fatigued tendon samples reached the secondary phase of fatigue loading, with a steady increase in the maximum cyclic displacement and cyclic stiffness following a rapid increase in the primary phase [63] (Figure 2). From cycle 1 to cycle 5000, the mean maximum cyclic displacement (*u*_*max*_) increased by 31% (4.10 to 5.35 mm) and cyclic stiffness (*k*) increased by 134% (140.45 N/mm to 328.66 N/mm). Detailed information of the fatigue behavior of the five fatigued samples is shown in the supplementary materials [56] (Figure S1).

**Figure 2.**
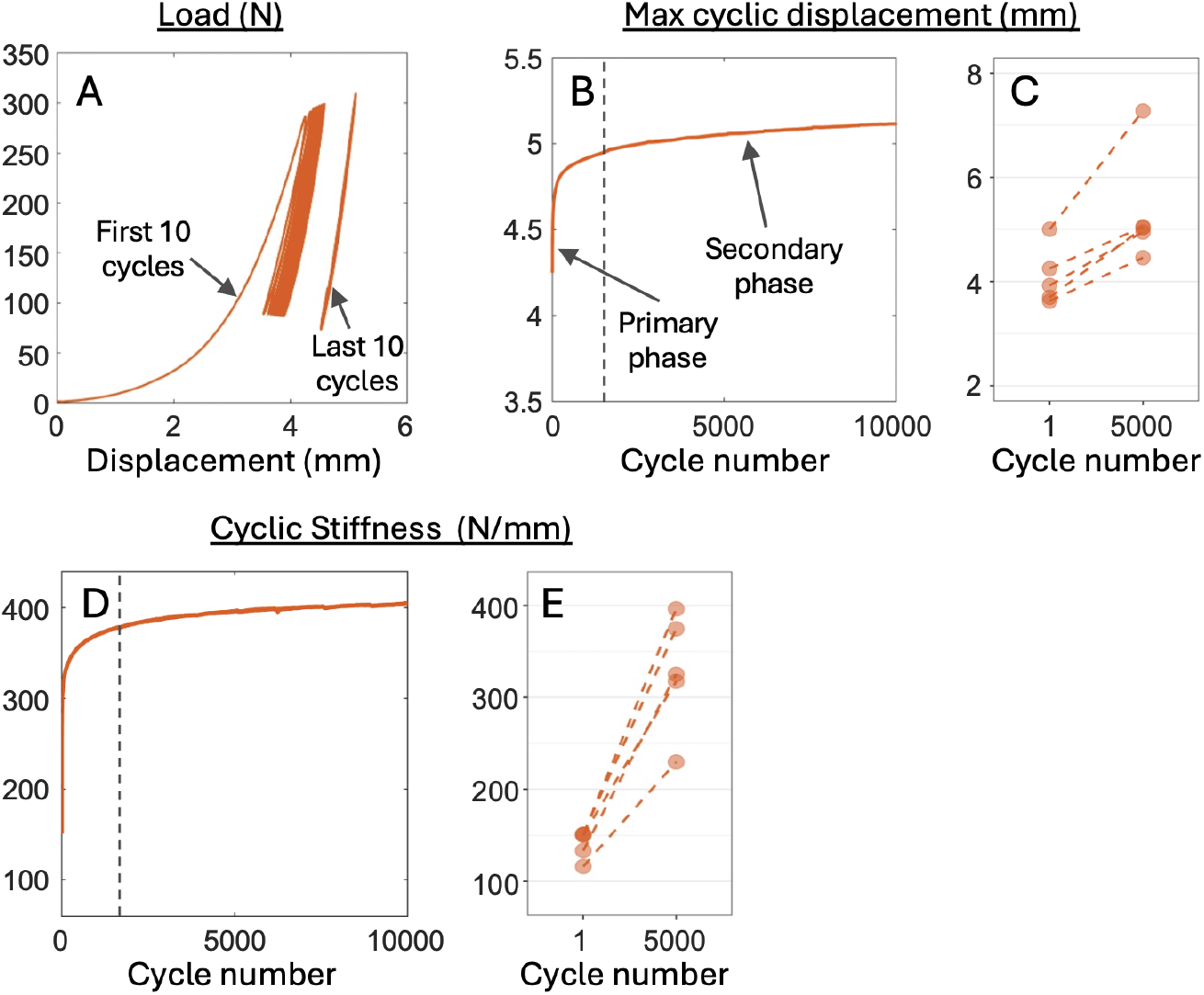
Fatigued tendons reached the secondary stage of fatigue loading prior to the DT-MRI assessment of fatigue-induced damage. The load-displacement curve from a representative fatigued tendon sample shows a creep behavior (A) that is characterized by a biphasic increase in maximum cyclic displacement (B). All fatigued tendons showed increase maximum cyclic displacement between cycle 1 and cycle 5000 of the fatigue protocol (C). An increase in cyclic stiffness was also measured in the representative tendon sample (D) and in all fatigued tendons between cycle 1 and cycle 5000 (E).

### 3.2. DT-MRI metrics are altered in microstructurally damaged fatigued tendons

To assess fatigue-induced changes in DT-MRI metrics, all tendons were scanned using a DT-MRI sequence before and after undergoing their respective protocols (Figure 1). The data collected from all the tendons (*n* = 10) were divided into nine anatomical regions to evaluate the spatial heterogeneity of the fatigue-induced damage (Figure 3A). The DT-MRI data of one sample from the control group was excluded due to vibration artifacts in the scanner bed. The results are presented as group medians, with differences in metrics referring to differences in group medians. Fatigue loading significantly increased Apparent Diffusion Coefficient (ADC) by 13% (*p* <0.001, Figure 3C), mainly attributable to a significant 13% increase in radial diffusivity (RD) (*p* <0.001, Figure 3D). Axial diffusivity (AD) moderately, but significantly, increased by 5% (*p* =0.027, Figure 3E). In contrast, all the DT-MRI metrics in the control group decreased; ADC and RD decreased by 16% (*p* <0.001), and AD by 14% (*p* <0.001). The ADC, RD, and AD of of the phosphate buffered saline surrounding the tendons did not significantly change between scans (*p >*0.42), demonstrating that any changes in diffusivity measured in the tendons were unlikely to be impacted by experimental conditions such as temperature changes between scans (Table S2). These data confirm the sensitivity of DT-MRI metrics to quantify the effects of mechanical fatigue in biological tissues.

**Figure 3.**
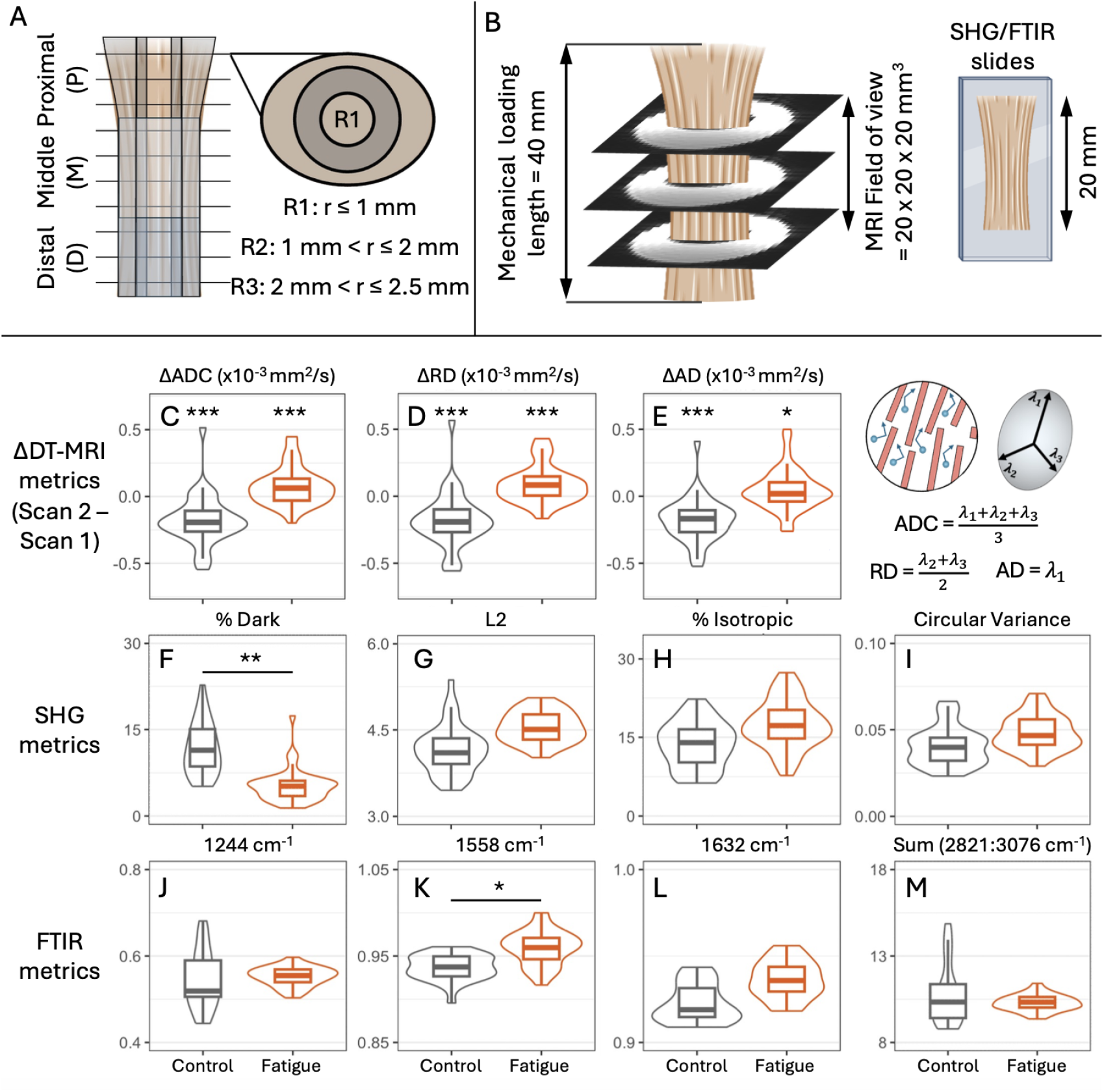
DT-MRI metrics, collagen fiber microstructure and secondary molecular structures are altered in fatigued tendons. (A) All imaging data were divided into nine anatomical regions to identify regional changes induced by fatigue loading. Each longitudinal subdivision (Proximal: P, Middle: M, Distal: D) was divided into three radial subdivisions (Central: R1, Intermediate: R2, Outer: R3). (B) MRI scans of the tendons were obtained in a centrally positioned field of view, with SHG and FTIR images collected from co-registered longitudinal slices. (C-E) Within-sample differences in DT-MRI metrics (measured between Scan 1 and Scan 2) show the different response of the tendon samples to the control and fatigue experiments. ADC, AD, and RD significantly increased in the fatigue group and significantly decreased in the control group. (F-I) SHG metrics measured in the control and fatigue group show between-group differences in the collagen fiber microstructure of the control and fatigue group, with fatigued samples having a significantly lower % Dark than the control samples. (J-M) FTIR metrics measured in the control and fatigue group showed a between-group change in the Amide II secondary protein structures (K), but did not cleave the covalent bonds in collagen (J), *β*-sheet (L), and C-H stretching nodes (M). The plots show the pooled regional data collected from all the samples. ^*^*p* <0.05, ^**^*p* <0.01, ^***^*p* <0.001

To probe whether collagen fiber disruption was a contributing mechanism underlying the changes in DT-MRI metrics, we co-registered and analyzed second harmonic generation (SHG) images of fatigued and control tendons (Figure 3B). SHG metrics confirmed the fatigue-induced changes in the collagen fiber microstructure of the tendons. Quantified as percent dark (% Dark), the inter-fibrillar spacing between collagen fibers and/or the number of fibers perpendicular to the imaging plane decreased from 11.4% in the control group to 5.2% in the fatigue group (*p* =0.002, Figure 3F). L2 is an SHG metric that quantifies the organization of the collagen fibers perpendicular to their principal orientation, approaching zero for highly aligned fibers. L2 non-significantly increased by 10% in the fatigue group (*p* =0.054, Figure 3G). Additionally, % Isotropic was measured as the percentage of areas in tendon with an isotropic structure. The control group had a % Isotropic of 13.99%, while the fatigue group had a % Isotropic of 17.24% (*p* =0.137, Figure 3H). Lastly, Circular Variance (CV) quantified the alignment of collagen fibers in the tendons, with a CV of 0 representing complete fiber alignment and a CV of 1 representing disorganized fibers. CV increased non-significantly by 17% in the fatigue tendons when compared to the controls (*p* =0.183, Figure 3I).

We conducted FTIR microscopy to study the molecular damage induced in the tendons by the fatigue loading protocol. The Amide I normalized absorbance differed significantly between fatigued and control samples at 1558 cm^*−*1^, Amide II, (*p* = 0.048, Figure 3K), indicating changes in secondary structures caused by the energy input of the fatigue loading protocol, breaking intermolecular forces and allowing the secondary protein structure to change. However, absorbance did not differ significantly between fatigued and control samples at 1244 cm^*−*1^ (collagen), 1632 cm^*−*1^ (*β*-sheet), or the sum of 2821-3076 cm^*−*1^ (C-H stretching modes), although an increasing trend is observed at 1632 cm^*−*1^ (Figure 3L). Additionally, the fatigue group showed less variation in absorbance than the control group at 1244 cm^*−*1^ and the sum of 2821:3076 cm^*−*1^ (Figure 3J,M).

### 3.3. Fatigue-induced changes in DT-MRI metrics are localized in regions with damaged collagen fiber microstructure

To assess the spatial heterogeneity of the fatigue-induced damage in the tendons, we evaluated the within-sample changes (difference between Scan 1 and Scan 2) in the DT-MRI metrics of nine anatomical regions (Figure 4A) and showed localized fatigue-induced damage in the proximal regions of the tendons. The proximal R1 region of the tendons in the fatigue group had a post-fatigue increase of 30% in ADC (*p* =0.001), 31% in RD (*p* =0.002), and 19% in AD (*p* =0.010). ADC increased by 16% ADC (*p* =0.017) and RD by 15% RD (*p* =0.004) in the proximal R2 region of the fatigued tendons (Figure 4B). No significant changes in DT-MRI metrics were measured in the remaining regions of the fatigued tendons, with modest increases in the middle regions and decreases in the distal regions. In the control group, only the distal intermediate region saw significant decreases in ADC (*p* <0.001), RD (*p* <0.001), and AD (*p* =0.003), while the remaining regions of the control tendons saw a non-significant decreasing trend. Changes in DT-MRI metrics in all the regions of the tendons (Figure S7) and statistics on the regional DT-MRI metrics of fatigued and control tendons (Tables S3-S5) are shown in the supplementary materials.

**Figure 4.**
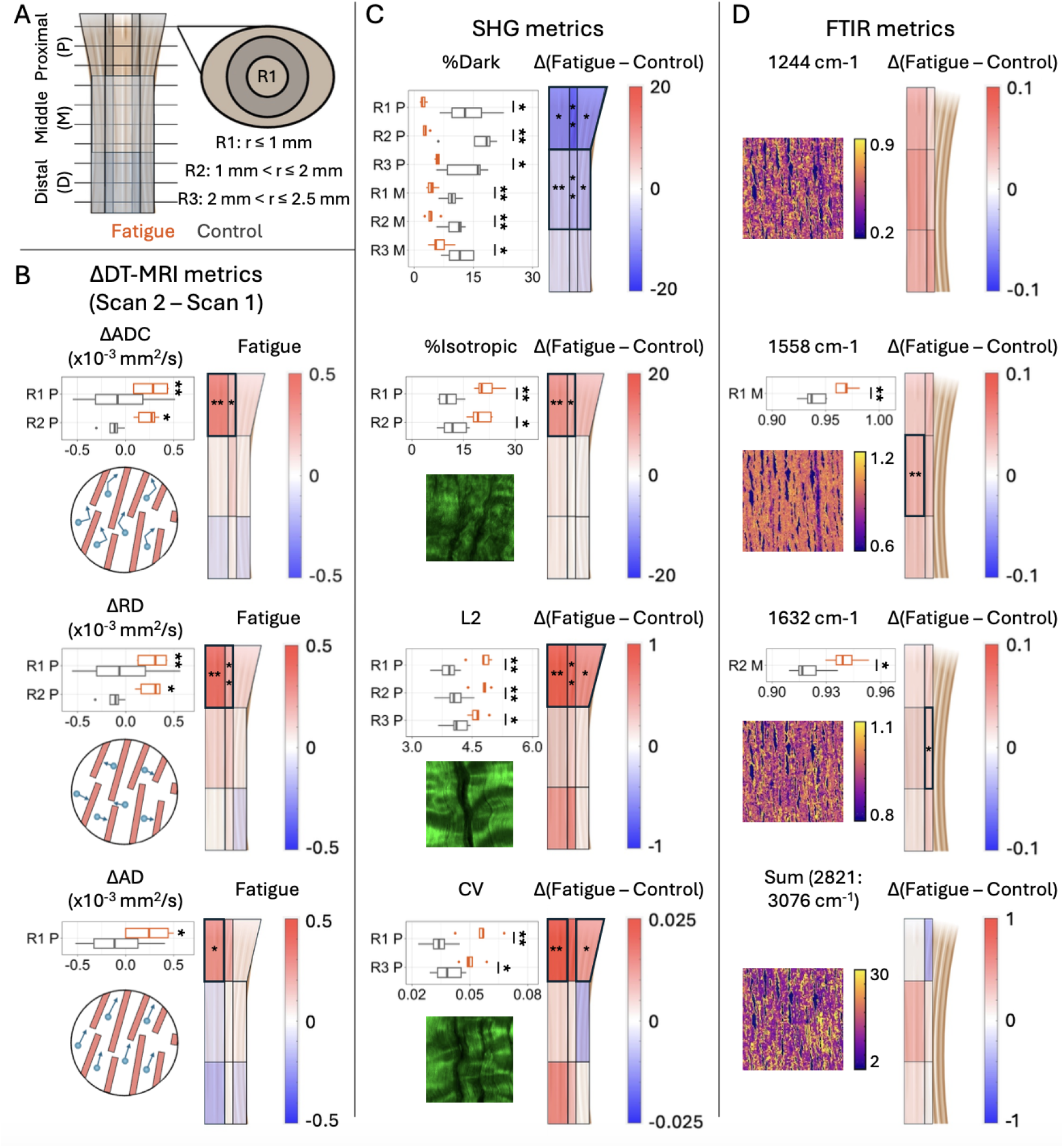
Fatigue-induced regional changes in DT-MRI metrics correspond to areas with damaged collagen fiber microstructure. (A) Tendon imaging data were divided into nine different anatomical regions to identify regional changes caused by fatigue loading. (B) within-sample changes in DT-MRI metrics (differences between Scan 1 and Scan 2) were region dependent in both the fatigue and control tendons. Boxplots show the regions with significant fatigue-induced increases from MRI Scan 1 to MRI Scan 2. The red-blue color maps show the change in the median regional DT-MRI metrics between Scan 1 and Scan 2 in the fatigue group. Statistical significance denotes within-sample differences in the DT-MRI metrics of the fatigue group. A schematic is shown to represent the directions of water molecular motion measured by each DT-MRI metric. (C) Between-group differences in the collagen fiber microstructure of the fatigue and control tendons via SHG. The boxplots show the data from the regions with significant changes in SHG metrics between the fatigue and control group. The red-blue color maps show the difference in the median regional SHG metrics between the fatigue and control group. The SHG images shown represent a region characterized by high % Isotropic, L2, and CV; field of view = 0.25 × 0.25 mm^2^. (D) Between-group differences in the molecular structures of the fatigue and control tendons via FTIR. The boxplots show the data from the regions with significant changes in FTIR metrics between the fatigue and control group. The red-blue color maps show the difference in the median regional FTIR metrics between the fatigue and control group. The rectangular images show the spatial variation in FTIR metrics in the middle R1 region of one fatigued sample; field of view = 2 × 2 mm^2^. Statistical significance in panels C and D denotes a between-group (fatigue vs. control) difference in the regional metrics. ^*^*p* <0.05, ^**^*p* <0.01, ^***^*p* <0.001

We confirmed the fatigue-induced changes in regional DT-MRI metrics with differences between the regional SHG metrics of the fatigued and control tendons. In the proximal R1 region, % Dark significantly decreased by 11% (*p* =0.045), % Isotropic increased by 10% (*p* =0.004), L2 increased by 22% (*p* =0.005) and CV increased by 67% (*p* =0.009) in the fatigue group when compared to the control group. We also measured a significant decrease of 15% in % Dark (*p* =0.007), an increase of 7% in % Isotropic (*p* =0.012), and an 18% increase in L2 (*p* =0.009) in the proximal R2 region of the fatigue group. In the proximal R3 region of the fatigue group, we measured a significant 10% decrease in % Dark significantly (*p* =0.048), as well as a 13% increase in L2 (*p* =0.023), and a 29% increase in CV (*p* =0.030). We also observed significant increases in the % Dark of the middle regions of the fatigued tendons when compared to the controls (Figure 4C). SHG metrics in all of the regions of the tendons are shown in the supplementary materials (Figure S7).

We also evaluated differences in regional FTIR metrics between the fatigued and control tendons to quantify regional molecular differences. Similar to the overall control and fatigue metrics, the regional FTIR imaging results indicate no significant changes in collagen and C-H absorbance in the fatigued tendons. The Amide I normalized Amide II absorbance (1558 cm^*−*1^) significantly increased by 3% in the medial R1 region (*p* = 0.003) of the fatigued tendons when compared to the control group, while the *β*-sheet absorbance (1632 cm^*−*1^) significantly increased by 2% in the medial R2 region (*p* = 0.012), as shown in Figure 4. No significant changes in any of the absorption bands were observed in the proximal and distal regions of the tendons. FTIR metrics in all the regions where data was collected are are shown in the supplementary materials (Figure S7).

### 3.4. Regional variation in fatigue-induced changes of DT-MRI metrics can be explained by underlying damage to collagen fiber microstructure

We investigated how regional changes in DT-MRI metrics can be explained by the variation in collagen fiber microstructure (regional SHG metrics) within the tendons by conducting repeated measures correlations. We found that regions with fatigue-induced increases in ADC were associated with less interfibrillar spacing and more fibers aligned in the imaging plane (% Dark, *r* =-0.63, *p* <0.001, Figure 5D), higher proportion of isotropically organized fibers (% Isotropic, *r* =0.32, *p* =0.045, Figure 5E), and lower transverse organization in collagen fibers (L2, *r* =0.34, *p* =0.032, Figure 5F). Fatigue-induced increases in RD were associated with lower % Dark (*r* =-0.67, *p* <0.001, Figure 5G) and higher L2 (*r* =0.36, *p* =0.021, Figure 5I), while fatigue-induced increases in AD were associated with lower % Dark (*r* =-0.51, *p* <0.001, Figure 5J) and higher % Isotropic (*r* =0.32, *p* =0.045, Figure 5K). No significant correlations were found between the changes in regional DT-MRI metrics and the collagen fiber microstructure of the tendons in the control group (Figure S8).

**Figure 5.**
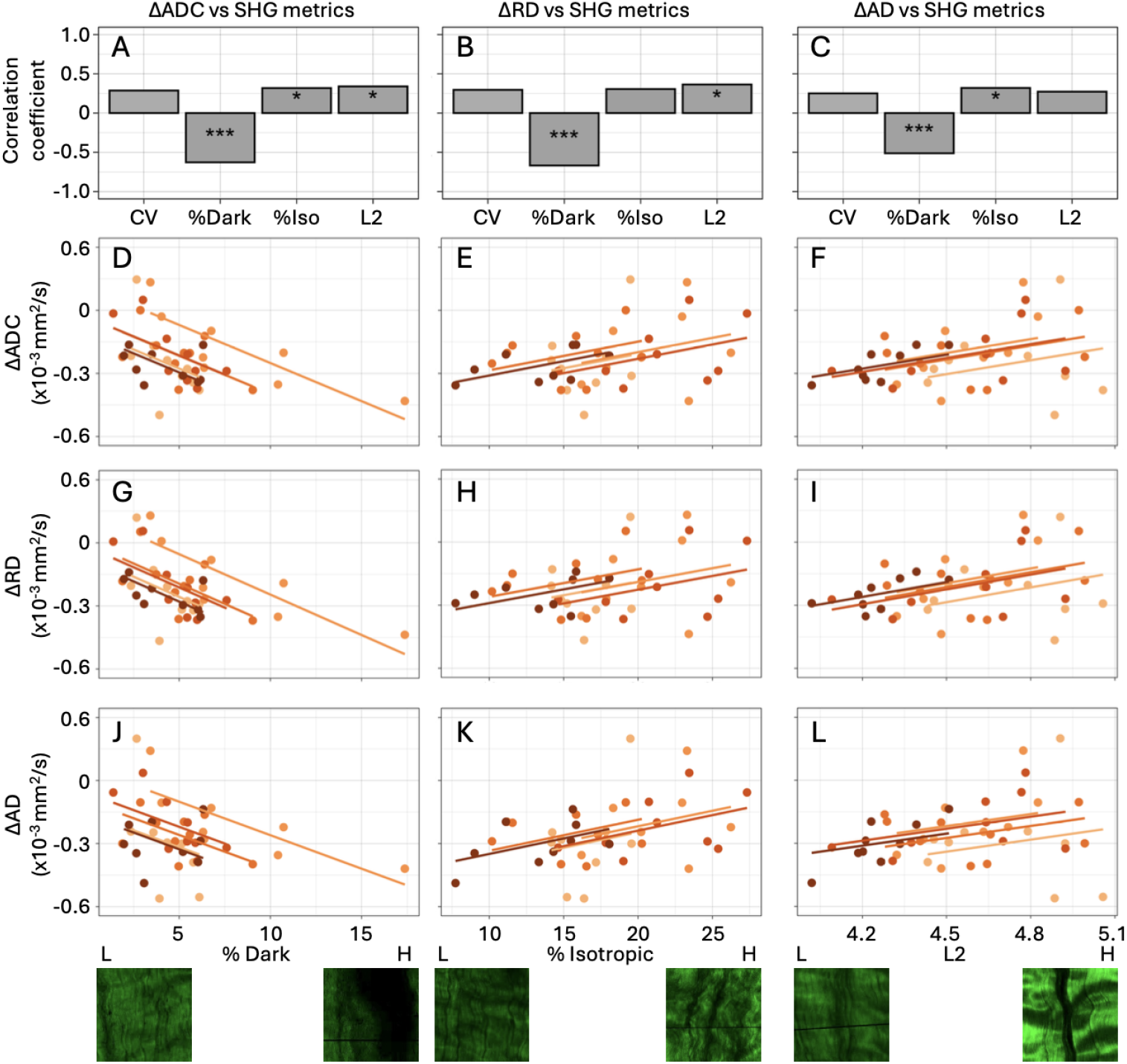
Variations in regional fatigue-induced changes in DT-MRI metrics depend on the damage to the region’s underlying collagen fiber microstructure. A-C) The variability in the regional fatigue-induced changes in the DT-MRI metrics can be explained by the SHG metrics describing the collagen fiber microstructure. Barplots show the correlation coefficients obtained from repeated measures correlations between the changes in DT-MRI metrics (Scan 2*−*Scan 1) and the SHG metrics of the fatigue group. Statistically significant correlations are labeled with ^*^*p* <0.05, ^**^*p* <0.01, ^***^*p* <0.001. (D-L) Repeated measures correlation plots showing the association between the changes in regional DT-MRI metrics and SHG metrics of all the tendons from the fatigue group. Each point represents one region of each sample, each line and shade of orange represents a different sample. Representative SHG images show the low (L) and high (H) values for the metrics considered. (Field of view = 0.25 × 0.25 mm^2^).

## 4. Discussion

This study identified how changes in DT-MRI metrics are associated with fatigue-induced microstructural and molecular damage in tendon. Significant increases in ADC, AD, and RD were observed after fatigue loading, in agreement with our findings in tendon-mimicking phantoms [48], confirming that water diffusivity increases in mechanically damaged fiber-based materials. The changes in DT-MRI metrics were localized in the proximal side of the tendons, where regional differences between the SHG metrics of the fatigued and control tendons were also observed. This region-specific response of the tendons to mechanical loading confirms the heterogeneity in the mechanical and microstructural properties of tendon, as noted in previous studies [51, 22]. While molecular (FTIR) changes in fatigued tendons were spatially unrelated to changes in DT-MRI metrics, the spatial correspondence of fatigue-induced changes in both DT-MRI and SHG metrics demonstrates that microstructural damage to the tendon’s collagen fiber microstructure results in increased water diffusivity within the tissue, even when there is no damage at the molecular level.

The significant correlations between the regional changes in DT-MRI and SHG metrics in the fatigued samples highlight the underlying relationships between tendon’s collagen fiber microstructure and DT-MRI metrics. We found that regional increases in ADC, AD, and RD were associated with a lower % Dark in the tendon regions. Since high % Dark is associated with the spacing between collagen fibers and a higher presence of out-of-plane fibers, the mechanically-damaged tendon regions with higher increases in diffusivity and lower % Dark were characterized by more densely packed and/or less crimped collagen fibers. This agrees with our previous findings in tendon-mimicking phantoms, were fatigue loading induced higher fiber compaction [48], and with previous results by Fung et al., where collagen fibers are initially uncrimped due to fatigue loading and damage increases progressively in localized areas of the tissue [24]. Tendons with fatigue-induced damage also have a higher presence of disorganized and kinked collagen fibers [24, 51], and this is supported by the significant correlation between the changes in ADC and AD with % Isotropic, showcasing how areas with a higher percentage of disorganized and kinked fibers were associated with increased diffusivity in the axial direction. Regional increases in ADC and RD were also correlated to increases in L2 in the tendons, resulting from the fatigue-induced disruption of collagen fibers perpendicular to the main axis of the tissue.

Using DT-MRI, the spatial heterogeneity in the response of the tendons to fatigue loading was non-invasively characterized across the whole tissue volume, in contrast to current histology methods that are limited to invasive assessments in localized fields of view [8, 24, 25]. Since the response of tendon to fatigue loading is spatially heterogeneous [22, 23], our work shows how a non-invasive tool can be used to detect local microstructural damages within a field of view that spans the entire tissue’s volume. This ability is of special interest in the understanding and diagnosis of soft tissue injuries [9, 10], where small areas of damage need to be identified within the whole tissue volume.

Molecular changes induced by fatigue in tendons were characterized with FTIR microscopy, showing that the levels of mechanical fatigue utilized in this study were insufficient to alter the structure of collagen molecules and C-H bonds. These results show that the changes in DT-MRI metrics observed in this study were caused by disruptions at the fiber microstructural level. This is in agreement with a previous study where collagen molecular damage was not ubiquitously observed after fatigue loading in tendons [23], whereas fatigue-induced disruptions to fiber microstructure are commonly observed [8, 24, 25, 23]. We did observe a regional increase in the *β*-sheet and Amide II content caused by the energy input of the fatigue loading protocol, but these occurred in medial regions of the tendons where DT-MRI changes were not observed. We also observed a narrowing in the absorbance distribution of collagen and C-H bonds in the fatigued tendons, which may indicate underlying changes in the *α* helix protein secondary structure absorption mode that the data was normalized to, but further study would be required to determine if this effect is real or meaningful.

In a preliminary study assessing the Achilles tendon in healthy participants and tendinopathy patients, ADC, AD, and RD were significantly lower in diseased tendons [42], suggesting that lower collagen content due to tendinopathy may be associated with lower ADC, AD, and RD. These results were also observed in experiments conducted in human Achilles tendons *in vitro*, were higher collagen content was associated with higher ADC, AD, and RD [64]. However, these studies are only representative of the baseline characteristics of the tendons at the time of a single DT-MRI scan, and did not assess progressive pathological or mechanically-induced changes within the same tendons at different time points. Additionally, elastin and cellular content mediate the DT-MRI metrics of soft tissue [65], and could confound changes in DT-MRI metrics observed in different study cohorts. Future studies could focus on elucidating the microstructural determinants of water diffusion in tendon, and how they could affect the outcomes of within-subject and between-subject studies.

The results from this study show promise to translate the use of DT-MRI as a soft tissue damage detection tool into the clinical setting. Longitudinal studies by van Dyck et al. measured DT-MRI metrics in patients who underwent ligament graft replacement [44] and internal bracing [45], and showed that DT-MRI metrics change during post-surgical recovery. Interestingly, van Dyck et al. suggested that changes in AD and RD could provide insights into the changes in ligament microstructure after remodeling [45]. Based on our results, ADC, AD, and RD could be used to track changes in the integrity of collagen fiber microstructure after clinical interventions, and to identify progressive tissue damage in populations at high risk of musculoskeletal injury.

Our study has several limitations. This study was conducted using tendon samples *in vitro*, so it did not represent physiological loading conditions experienced by tendons *in vivo*. The cellular remodeling that occurs in tendons *in vivo* can not be accounted for in our study, and its effect in the DT-MRI metrics measured after physiological loading in tendon remains unknown. The use of *in vitro* loading also limited the variability seen with FTIR imaging as inflammation [66] and biochemical modifications were not possible. Previous studies in deep digital flexor tendons have shown that the section of the deep digital flexor we studied, distal to the tendon’s bifurcation, have complex collagen microstructure patterns due to a combination of tensile and compressive physiological loads applied to the tendon [67]. Because we applied uniaxial tensile loading to the tendon’s samples, the fatigue-induced microstructural changes measured in this study do not replicate the changes expected in physiological conditions. However, the damage created by our loading protocol allowed us to evaluate the sensitivity of DT-MRI metrics to fatigue-induced damage in tendon. Finally, we did not evaluate the potential effect of fatigue-induced changes in the elastin, proteoglycan, and cellular components of the tendons, which could be evaluated in future work.

Having shown that DT-MRI metrics are sensitive to fatigue-induced damage to tissue microstructure in tendon *in vivo*, future studies should focus on using DT-MRI metrics as potential biomarkers to non-invasively assess tendon health *in vivo*. Studies of physiological tendon fatigue loading in animal models and human participants should be conducted to confirm our findings and relate changes in DT-MRI metrics to clinical assessments of tissue health, such as joint passive laxity and active range of motion. Moreover, detailed assessments of tendon fiber architecture could be done using DT-MRI based fiber tractography. Finally, relating changes in DT-MRI metrics to the mechanical properties of tendon also needs future study, as it has been found in arteries that DT-MRI data can be related to the mechanical properties of collagen-based tissues [68].

This study shows that changes in DT-MRI metrics are capable of detecting mechanically induced microstructural damage in tendons *in vitro*. We show that local fatigue-induced changes in ADC, AD, and RD correspond to local areas of fatigue-induced damage in the collagen fiber microstructure of tendon, showcasing the ability of DT-MRI to non-invasively detect localized microstructural damage. We found significant correlations between changes in DT-MRI metrics and the underlying collagen fiber microstructure metrics of fatigued tendons, showing that changes in DT-MRI metrics can be explained by microstructural disruptions observed in tendon after fatigue loading. The minimal variation in IR absorption bands showed that the microstructural changes determined with DT-MRI and SHG were not influenced by changes to the molecular makeup of the tissue. In conclusion, our results show that DT-MRI is a non-invasive and potentially translatable tool capable of clinically detecting mechanopathologies in tendon, and possibly other connective soft tissues.

## 5. Acknowledgements

We are thankful to Mahmuda Arshee, Kellie Halloran, Griffin Sipes, Melany Opolz, Michael Focht, Mikayla Hoyle, and Elahe Ganji for their assistance with the experiments. This work was conducted in part at the Biomedical Imaging Center of the Beckman Institute for Advanced Science and Technology at the University of Illinois Urbana-Champaign (UIUC-BI-BIC). DT-MRI scans were funded by a seed grant from UIUC-BI-BIC. BMD was funded by National Institutes of Health grant R01 AR073831. MPC acknowledges partial support from a National Science Foundation (NSF) Division of Ocean Sciences Postdoctoral Fellowship (2205819).

