## Supplementary materials for "Non-invasive detection of local microstructural damage in tendon via Diffusion Tensor MRI"

### Supplementary Materials for
### Non-invasive detection of local microstructural damage in tendon via Diffusion Tensor MRI

Roberto A. Pineda Guzman, Amir Ostadi Moghaddam, Matthew P. Confer,
Shreyan Majumdar, Rohit Bhargava, Amy J. Wagoner Johnson, Bruce M. Damon,
Mariana E. Kersh*

#### Materials and Methods

##### Tendon preparation prior to imaging and image co-registration

Prior to the DT-MRI scans, the tendons were placed inside a 15 mL centrifuge tube filled with 1x phosphate buffered saline (PBS) and pre-stretched to remove slack and standardize the tendon's reference length. Both ends of the tendons were attached to a string using cyanoacrylate glue. The string on the distal end of the tendon was attached to the centrifuge tube, while the string on the proximal end was loaded with a hanging 0.98 N aluminum weight, pre-stretching the tendon (Figure S2A). The string connecting the tendon to the aluminum weight was then fixed to the tube's cap to fix the tendon's pre-stretched configuration (Figure S2B). A registration marker filled with MR-visible fluid was used to ensure that the MRI scan's field of view would start in the proximal edge of the tissue dye mark.

After fatigue loading, the tendon was pre-stretched using the same protocol for the pre-fatigue scan (Figure S2C). Using the tissue dye mark placed on the tendon, the field of view of the post-fatigue scan was located in the same central area scanned pre-fatigue using a registration marker (Figure S2D). This step helped ensure that the same area of the tendon was assessed in the pre and post-fatigue DT-MRI scans. For the tendons in the control group, the sample was prepared as shown in Figure S2A-B, and the registration marker remained in the same place between the Day 1 and Day 2 scans to help ensure that the same section of the tendon was scanned.

#### Evaluation of DT-MRI analyses using signal averaging

To evaluate whether the signal averaging method used to process the Diffusion Tensor Imaging data does not skew the original image statistics and effectively reduces the effect of noise, we ran a Monte Carlo simulation scheme in MATLAB (version R2023a, Mathworks, Natick, MA, USA) - similar to previously published work by Froeling et al. (*37*), Damon (*36*), and Anderson (*35*). We defined 2 ROIs for each of the regions evaluated in the tendons, with variable voxel size depending on the size of the regions, and random and normally distributed signal intensities, $S_0$. Each voxel was assigned a diffusion tensor, $\mathbf{D}$, with random and normally distributed axial diffusivity (AD), radial diffusivity (RD), and a principal eigenvector oriented at an angle with respect to the tendon's longitudinal axis, $\theta$. The data distributions were generated using the *normrnd* function. The mean and standard deviation of the parameters input into the function are listed in Table S1. The mean and standard deviation of the SNR used in the simulation were chosen based on the region with lowest SNR (i.e. R1-Min) of the region and the region whose SNR was closest to the mean SNR measured on all the radial region's metrics (i.e. R1-Mean). The mean AD and RD of each region were chosen based on the mean AD and RD metrics of each radial region, and the standard deviation was chosen to be 10% of the mean data. A mean $\theta = 0$ was chosen assuming alignment of the tendon's diffusion direction along the tissue's main axis, with an angular standard deviation of 10 degrees across both the transverse plane of the tissue. An example of the simulated and randomly generated ROI used in this Monte Carlo scheme is shown in Figure S3. Fractional anisotropy (FA) was computed from AD ($\lambda_1$) and RD, assuming $RD = \lambda_2 = \lambda_3$ as shown in equation S1.

$$FA = \sqrt{\frac{(\lambda_1 - \lambda_2)^2 + (\lambda_2 - \lambda_3)^2 + (\lambda_3 - \lambda_1)^2}{2(\lambda_1^2 + \lambda_2^2 + \lambda_3^2)}} \tag{S1}$$

The diffusion weighted signals of the ROIs were computed using the same b-value and diffusion directions, $\mathbf{g}$, specified in the DT-MRI experiment (Equation S2). Random noise with unit variance was added to the ROI signals using MATLAB's *randn* function.

$$S_{DW} = S_0 e^{-b\mathbf{g}\mathbf{D}\mathbf{g}^T} \tag{S2}$$

The ADC, AD, RD, FA and $\theta$ of each noisy ROI were computed by averaging the signals in the ROI, obtaining a diffusion tensor and its respective DT-MRI metrics from each of the 6 ROIs generated. The simulation was repeated 5000 times. The computed ADC, AD, RD, FA, and Z angle from the noisy ROIs were compared to the prescribed DT-MRI metrics of each ROI.

Figures S4-S6 show the results of the simulations for each respective ROI. The prescribed DT-MRI metrics of each ROI are shown with the gray dashed line, and intervals specifying ±5% difference with respect to the prescribed value are shown with the gray dotted lines on the plots for ADC, AD, and RD. It can be seen from the plots below that ADC, AD, and RD are consistently computed within a 5% difference of the prescribed values for all the ROIs, even at the lowest SNR level specified in the region R1. However, FA is consistently underestimated on regions R2 and R3. This could be due to the signal averaging process, as it has been seen by others that it could result in decreased FA, especially when there is heterogeneity introduced in the primary diffusion direction [4]. $\theta$ is consistently overestimated in all the ROIs, mainly caused by the low SNR levels measured in the tendon and implemented in this simulation. These results show that at the conditions in which the tendons were imaged in our study, ADC, AD, and RD are DT-MRI metrics that can be measured with almost no bias and ±5% precision. However, FA and $\theta$ cannot be accurately measured under the tendon's imaging conditions.

#### Supplementary Results

##### Fatigue loading of tendons

The mechanical behavior all the fatigued tendons is shown in FigureS1. Samples 1 and 2 completed the 10000 loading cycles of the loading protocol with no interruption. Sample 3 slipped in the grips during the initial 3089 loading cycles. It was re-gripped and underwent 7500 additional loading cycles, for a total of 10589 loading cycles. Sample 4 had its fatigue protocol interrupted during cycle 7500 but completed the fatigue loading protocol with 2500 additional loading cycles, for a total of 10000 loading cycles. Sample 5 underwent 5766 loading cycles before abruptly failing on its distal end and could not be re-gripped. Sample 5 was used in the study because the area of the tendon imaged in the DT-MRI scans showed no macroscopic signs of damage.

**Changes in ADC, AD, and RD of the PBS surrounding the tendons in fatigue and control DT-MRI scans**

The ADC, AD, and RD of the Phosphate Buferred Saline (PBS) surrounding the tendons did not change between Scan 1 and Scan 2 of the fatigue and control group. After checking for normality using a Shapiro-Wilk test, we conducted t-tests and found no significant differences in ADC, AD, and RD between the Scan 1 and 2 of the fatigue and control groups scans (Table S2).

**Correlations between changes in ADC, AD, and RD and the SHG metrics measured in the control group**

The correlation coefficients between the changes in ADC, AD, and RD and the SHG metrics measured in the tendons of the control group are shown in Figure S8. No significant correlations were found in the control group data.

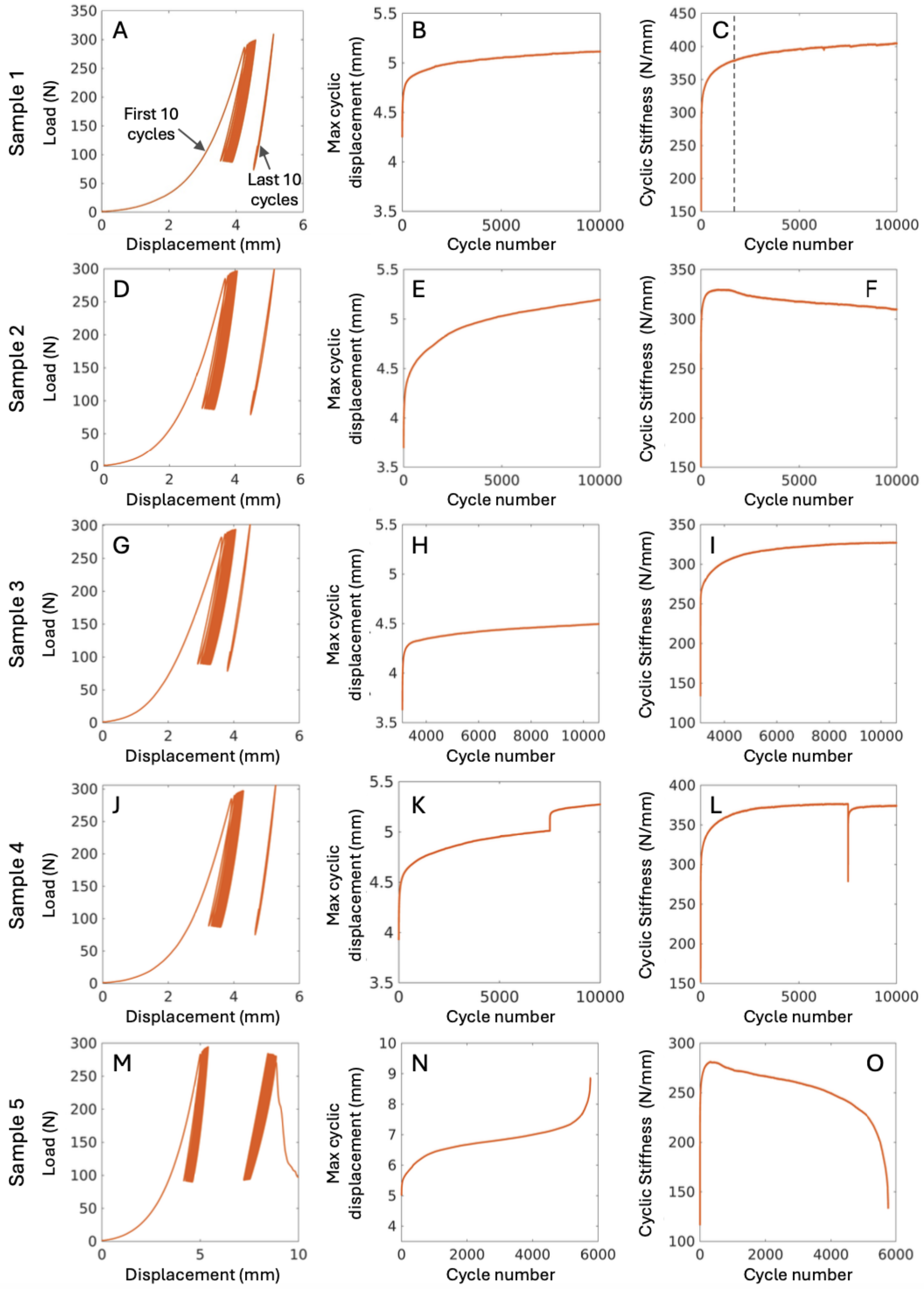


**Figure S1**: Plots showing the load-displacement curve, maximum cyclic displacement, and cyclic stiffness of the five fatigued tendons. Each row contains the fatigue loading data from each tendon sample.

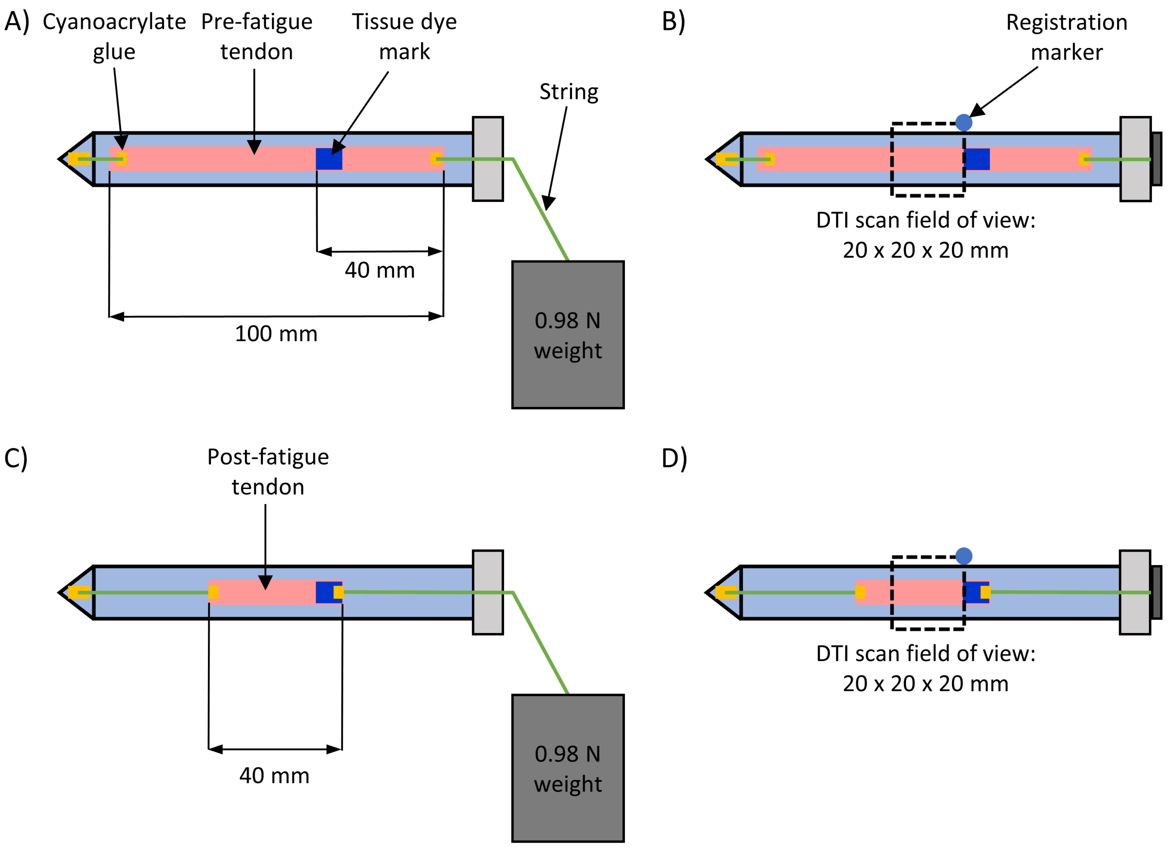


**Figure S2**: A) Prior to MRI Scan 1, the tendons were pre-stretched and a tissue dye mark was applied to the tendon to help ensure that the central area was imaged. B) A registration marker was placed over the tissue dye to guide the positioning of the field of view in MRI Scan 1. C) The ends of the tendon were removed after the fatigue loading protocol and the tendon was pre-stretched prior to MRI Scan 2. D) The tissue dye mark was used to position the field of view in MRI Scan 2 and helped ensure that the same region of the tendon was imaged in both scans.

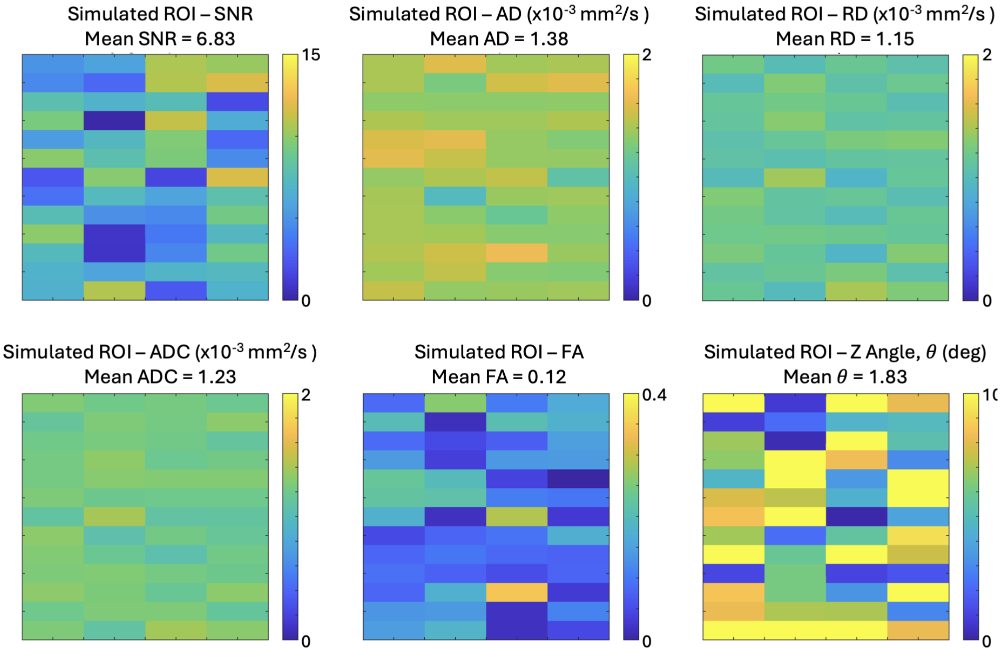


**Figure S3**: Prescribed SNR, ADC, AD, RD, and $\theta$ used to generate the noise-free diffusion weighted signals in the ROI that represented the minimum SNR measured in region R1.

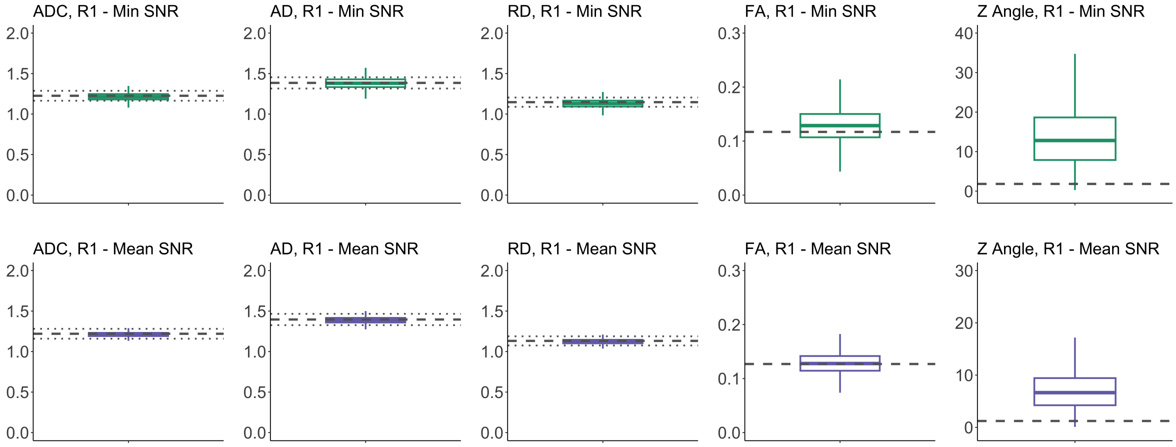


**Figure S4**: DT-MRI metrics measured on ROIs corresponding to region R1 of the tendons, with minimum SNR conditions (top row) and mean SNR conditions (bottom row). The boxplots represent the results of the signal averaging measuring method in the 5000 Monte Carlo trials, the dashed horizontal line represents the prescribed DT-MRI .

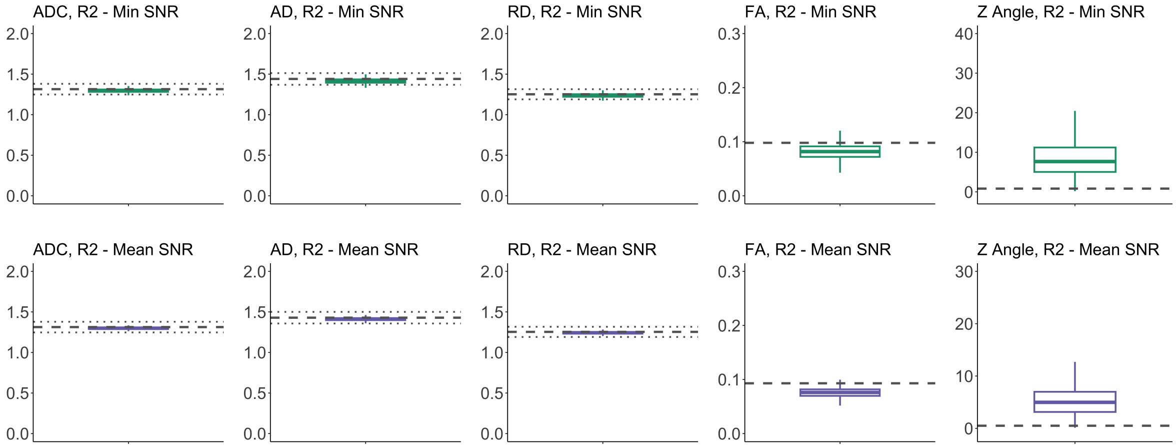


**Figure S5**: DT-MRI metrics measured on ROIs corresponding to region R2 of the tendons, with minimum SNR conditions (top row) and mean SNR conditions (bottom row). The boxplots represent the results of the signal averaging measuring method in the 5000 Monte Carlo trials, the dashed horizontal line represents the prescribed DT-MRI .

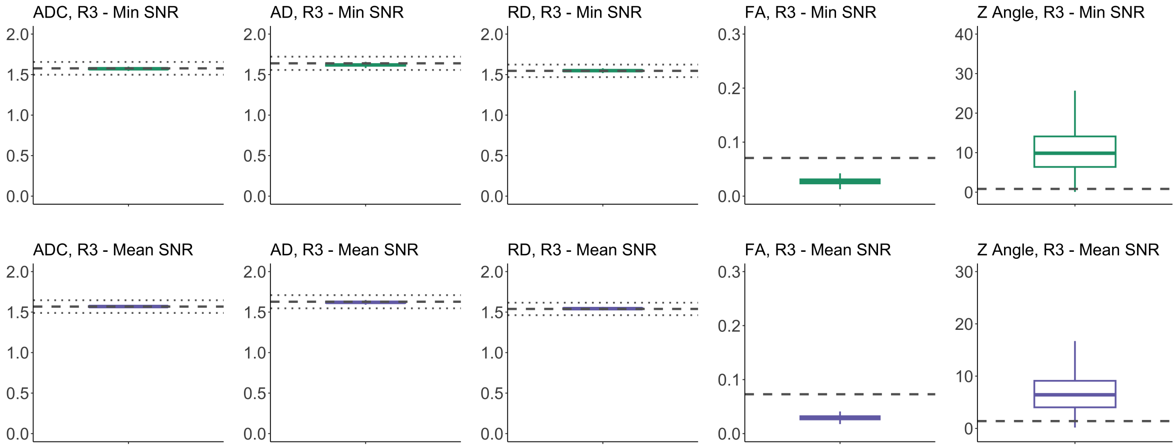


**Figure S6**: DT-MRI metrics measured on ROIs corresponding to region R3 of the tendons, with minimum SNR conditions (top row) and mean SNR conditions (bottom row). The boxplots represent the results of the signal averaging measuring method in the 5000 Monte Carlo trials, the dashed horizontal line represents the prescribed DT-MRI .

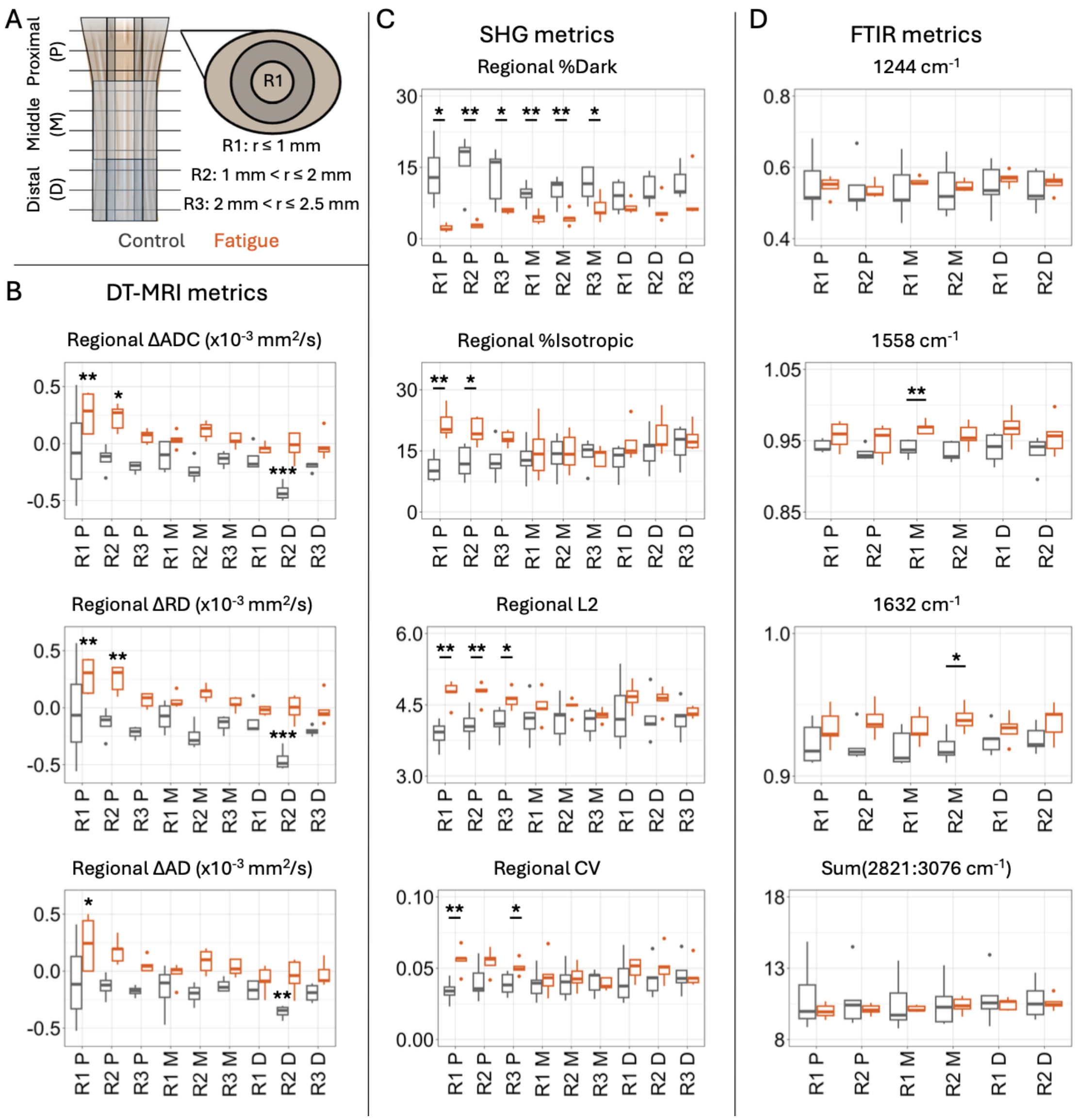


**Figure S7**: (A) Tendon imaging data were divided into nine different anatomical regions to identify regional changes caused by fatigue loading. Region R1 corresponds to the central radial region, while R2 and R3 correspond to the intermediate and outer radial regions of the tendons. (B) Boxplots showing the fatigue-induced changes in the regional DT-MRI metrics of the tendons. (C) Boxplots showing the regional SHG metrics measured in the tendons from the fatigue and control group. (D) Boxplots showing the regional FTIR metrics measured in the tendons from the fatigue and control group. FTIR data was not available for region R3 of the tendons.*$p$ <0.05, **$p$ <0.01, ***$p$ <0.001

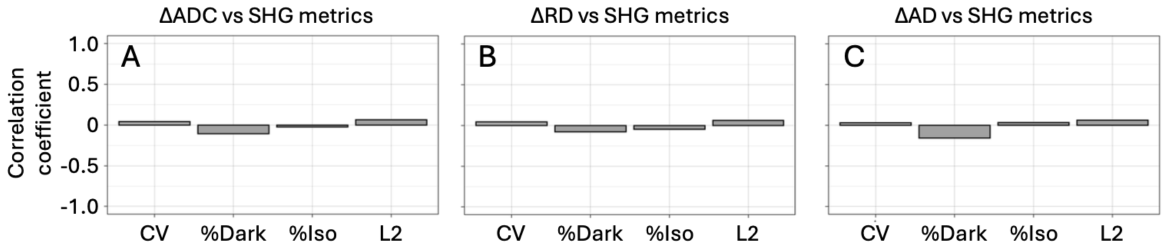


**Figure S8**: Correlation coefficients measured between the changes in ADC (A), AD(B), and RD (C) and the SHG metrics measured in the tendons of the control group.

**Table S1**: Mean ± standard deviation of the prescribed parameters in the simulated ROIs.

| | ROIs created | | | | | |
|---|---|---|---|---|---|---|
| ROI parameters | R1 - Min | R1 - Mean | R2 - Min | R2 - Mean | R3 - Min | R3 - Mean |
| SNR | 6.8 ± 4.0 | 13.2 ± 7.4 | 10.8 ± 8.6 | 17.8 ± 11.3 | 16.6 ± 14.3 | 25.3 ± 21.0 |
| AD (x$10^{-3}$ $mm^2/s$) | 1.37 ± 0.137 | 1.37 ± 0.137 | 1.43 ± 0.143 | 1.43 ± 0.143 | 1.64 ± 0.164 | 1.64 ± 0.164 |
| RD (x$10^{-3}$ $mm^2/s$) | 1.15 ± 0.115 | 1.15 ± 0.115 | 1.25 ± 0.125 | 1.25 ± 0.125 | 1.54 ± 0.154 | 1.54 ± 0.154 |
| Principal diffusion direction angle, $\theta$ (degrees) | 0 ± 10 | 0 ± 10 | 0 ± 10 | 0 ± 10 | 0 ± 10 | 0 ± 10 |
| Number of voxels in ROI | 52 | 42 | 111 | 111 | 235 | 183 |

**Table S2**: Mean ± standard deviation values of ADC, AD, and RD measured in the PBS surrounding the Scans 1 and 2 of the fatigue and control group.

| | Fatigue experiment | | Control experiment | |
|---|---|---|---|---|
| DT-MRI metric (1 x $10^{-3}$ $mm^2/s$) | Scan 1 | Scan 2 | Scan 1 | Scan 2 |
| ADC | 1.93 ± 0.03 | 1.93 ± 0.03 | 1.90 ± 0.06 | 1.87 ± 0.06 |
| RD | 1.90 ± 0.03 | 1.90 ± 0.03 | 1.88 ± 0.06 | 1.84 ± 0.06 |
| AD | 1.99 ± 0.04 | 1.99 ± 0.04 | 1.96 ± 0.06 | 1.92 ± 0.06 |

**Table S3**: Median ± median absolute deviation of the ADC ($1x10^{-3}$ $mm^2/s$) measured in the different regions of fatigued and control tendons.

| Region | | Fatigue experiment | | Control experiment | |
|---|---|---|---|---|---|
| | | Scan 1 | Scan 2 | Scan 1 | Scan 2 |
| Proximal | Central | 1.23 ± 0.04 | 1.6 ± 0.12 | 1.18 ± 0.41 | 1.11 ± 0.39 |
| | Intermediate | 1.30 ± 0.11 | 1.51 ± 0.05 | 1.40 ± 0.04 | 1.19 ± 0.17 |
| | Outer | 1.54 ± 0.05 | 1.63 ± 0.03 | 1.66 ± 0.02 | 1.43 ± 0.06 |
| Middle | Central | 1.31 ± 0.07 | 1.34 ± 0.17 | 1.30 ± 0.18 | 1.19 ± 0.05 |
| | Intermediate | 1.23 ± 0.08 | 1.35 ± 0.08 | 1.50 ± 0.03 | 1.20 ± 0.09 |
| | Outer | 1.66 ± 0.04 | 1.68 ± 0.06 | 1.65 ± 0.05 | 1.51 ± 0.09 |
| Distal | Central | 1.17 ± 0.13 | 1.09 ± 0.13 | 1.18 ± 0.08 | 1.05 ± 0.10 |
| | Intermediate | 1.27 ± 0.07 | 1.30 ± 0.05 | 1.64 ± 0.05 | 1.18 ± 0.05 |
| | Outer | 1.62 ± 0.02 | 1.54 ± 0.03 | 1.66 ± 0.02 | 1.48 ± 0.02 |

**Table S4**: Median ± median absolute deviation of the RD ($1x10^{-3}$ $mm^2/s$) measured in the different regions of fatigued and control tendons.

| Region | | Fatigue experiment | | Control experiment | |
|---|---|---|---|---|---|
| | | Scan 1 | Scan 2 | Scan 1 | Scan 2 |
| Proximal | Central | 1.19 ± 0.10 | 1.55 ± 0.09 | 1.08 ± 0.38 | 1.05 ± 0.45 |
| | Intermediate | 1.28 ± 0.14 | 1.47 ± 0.07 | 1.33 ± 0.06 | 1.12 ± 0.20 |
| | Outer | 1.50 ± 0.04 | 1.59 ± 0.03 | 1.63 ± 0.01 | 1.38 ± 0.07 |
| Middle | Central | 1.21 ± 0.12 | 1.30 ± 0.13 | 1.2 ± 0.13 | 1.09 ± 0.05 |
| | Intermediate | 1.19 ± 0.10 | 1.30 ± 0.07 | 1.44 ± 0.03 | 1.11 ± 0.10 |
| | Outer | 1.62 ± 0.04 | 1.65 ± 0.03 | 1.61 ± 0.08 | 1.46 ± 0.09 |
| Distal | Central | 1.04 ± 0.14 | 1.04 ± 0.11 | 1.08 ± 0.11 | 0.96 ± 0.12 |
| | Intermediate | 1.20 ± 0.06 | 1.23 ± 0.07 | 1.61 ± 0.04 | 1.10 ± 0.07 |
| | Outer | 1.58 ± 0.04 | 1.50 ± 0.06 | 1.62 ± 0.03 | 1.43 ± 0.04 |

**Table S5**: Median ± median absolute deviation of the AD (1x$10^{-3}$ $mm^2/s$) measured in the different regions of fatigued and control tendons.

| Region | | Fatigue experiment | | Control experiment | |
|---|---|---|---|---|---|
| | | Scan 1 | Scan 2 | Scan 1 | Scan 2 |
| Proximal | Central | 1.41 ± 0.14 | 1.68 ± 0.19 | 1.39 ± 0.40 | 1.26 ± 0.32 |
| | Intermediate | 1.41 ± 0.13 | 1.60 ± 0.08 | 1.51 ± 0.08 | 1.34 ± 0.07 |
| | Outer | 1.63 ± 0.10 | 1.69 ± 0.05 | 1.71 ± 0.05 | 1.53 ± 0.04 |
| Middle | Central | 1.5 ± 0.07 | 1.43 ± 0.14 | 1.45 ± 0.35 | 1.38 ± 0.03 |
| | Intermediate | 1.43 ± 0.17 | 1.44 ± 0.09 | 1.62 ± 0.05 | 1.38 ± 0.07 |
| | Outer | 1.72 ± 0.04 | 1.75 ± 0.11 | 1.74 ± 0.02 | 1.60 ± 0.09 |
| Distal | Central | 1.35 ± 0.15 | 1.19 ± 0.10 | 1.38 ± 0.07 | 1.23 ± 0.09 |
| | Intermediate | 1.41 ± 0.11 | 1.41 ± 0.06 | 1.70 ± 0.07 | 1.35 ± 0.02 |
| | Outer | 1.68 ± 0.01 | 1.6 ± 0.03 | 1.70 ± 0.08 | 1.57 ± 0.02 |
